# A comparative study of population genetic structure reveals patterns consistent with selection at functional microsatellites in common sunflower

**DOI:** 10.1101/2021.08.01.454655

**Authors:** Chathurani Ranathunge, Melody Chimahusky, Mark E Welch

**Affiliations:** Mississippi State University

**Keywords:** *Helianthus annuus*, microsatellites, selection, population genetics, STR, sunflower

## Abstract

Microsatellites have long been considered non-functional, neutrally evolving regions of the genome. Recent findings suggest that they can function as drivers of rapid adaptive evolution. Previous work on common sunflower identified 479 transcribed microsatellites where allele length significantly correlates with gene expression (eSTRs) in a stepwise manner. Here, a population genetic approach is used to test whether eSTR allele length variation is under selection. Genotypic variation among and within populations at 13 eSTRs was compared with that at 19 anonymous microsatellites in 672 individuals from 17 natural populations of sunflower from across a cline running from Saskatchewan to Oklahoma. Expected heterozygosity, allelic richness, and allelic diversity were significantly lower at eSTRs, a pattern consistent with higher relative rates of purifying selection. Further, an analysis of variation in microsatellite allele lengths (*lnRV*), and heterozygosities (*lnRH*), indicate recent selective sweeps at the eSTRs. Mean microsatellite allele lengths at four eSTRs within populations are significantly correlated with latitude consistent with the predictions of the tuning knob model which predicts stepwise relationships between microsatellite allele length and phenotypes. This finding suggests that shorter or longer alleles at eSTRs may be favored in climatic extremes. Collectively, our results imply that eSTRs are likely under selection and that they may be playing a role in facilitating local adaptation across a well-defined cline in the common sunflower.

## INTRODUCTION

Microsatellites, or short tandem repeats (STRs), are ubiquitous repetitive elements in prokaryotic and eukaryotic genomes. They typically consist of tandemly repeated motifs that are 1 – 6 bp in length. Microsatellites are known for their elevated mutation rates that are orders of magnitude greater than that of point mutations (Bhargava & Fuentes, 2010), and these high mutation rates may involve mechanisms such as replication slippage (Tautz & Renz, 1984). Because of the variability of microsatellites and the assumption that they are non-functional, and therefore evolve in a neutral fashion, microsatellites were once considered the “ molecular marker of choice” in population genetics and forensic analyses (Hodel *et al*., 2016; Oliveira *et al*., 2006). However, a growing body of research now implies that microsatellites can be functional and can be targets of natural selection. Microsatellites located within or close to genic regions are particularly important in this regard as changes in length of these microsatellites are likely to influence gene expression or protein structure (Li *et al*., 2004; Gemayel *et al*., 2010).

Initial evidence for a potential functional role for microsatellites comes from research that linked microsatellite expansion to human diseases such as Huntington’s disease (Andrew *et al*., 1993), Fragile X syndrome (Kremer *et al*., 1991), and Frederic’s ataxia (Dürr *et al*., 1996). Research also implicates microsatellites as a source of adaptation. Microsatellites have now been linked to reproductive success in some mammals (Lonn *et al*., 2017), neuronal and craniofocal development in primates (Ohadi *et al*., 2015), and several quantitative phenotypes in *Arabidopsis thaliana* strains (Press et al, 2018). It has been further proposed that microsatellites can function as “ tuning knobs” where stepwise changes in microsatellite allele length can result in stepwise changes in phenotypes (Kashi *et al*., 1997; King *et al*., 1997; Trifonov, 2004), allowing for fine-tuning of quantitative phenotypes in response to environmental stresses. These studies that highlight the adaptive potential of microsatellites call into question the assumption that microsatellites are non-functional and therefore neutrally evolving genomic regions. In the vast majority of microsatellite-based empirical studies, signatures of selection detected at microsatellites are typically ascribed to genetic hitchhiking (Schlötterer, 2002; Rockman *et al*., 2005; Schlötterer & Dieringer, 2005; Gross *et al*., 2007; Kane & Rieseberg, 2007, 2008). Only the rare study has explored the possibility that microsatellites themselves are targets of natural selection (Haasl & Payseur, 2013; Haasl *et al*., 2014, Lonn *et al*., 2017, Pramod *et al*., 2014). Work on the bank vole, *Myodes glareolus*, reveals evidence for stabilizing selection against long and short allele lengths in microsatellites located in the arginine vasopressin receptor 1a (avpr1a) and oxytocin receptor (oxtr) loci that play an evolutionary significant role in vertebrate social and sexual behavior (Watts *et al*., 2017). In another study on *Arabidopsis thaliana*, strong selection was implicated acting on a microsatellite located within the PHYTOCHROME AND FLOWERING TIME 1(PFT1) gene linked to regulation of flowering (Rival *et al*., 2014). If these anecdotal findings on microsatellites under selection are generalizable, it would provide evidence for a novel mechanism by which natural populations may rapidly acquire adaptive variation.

The adaptive potential of a locus is often exclusively inferred by polymorphism-based tests of selection or population genetic analyses. However, inferring the adaptive potential of a locus with population genetic analyses alone can be challenging. For example, false positive estimates of selection could be attributable to population structure dependent on the sampling scheme rather than positive selection itself (Slatkin & Wiehe, 1998). To better support inferences concerning adaptation at specific loci, Storz & Wheat (2010) suggest an integrative approach where population genetic analyses are combined with functional experiments. Such integrative studies have been shown effective in providing insights into potential adaptive mechanisms (Watt *et al*., 1996, 2003; Lamason *et al*., 2005; Linnen *et al*., 2009). In some cases, evidence of positive selection detected by population genetic analyses may motivate functional experiments to investigate whether candidate loci identified do in fact contribute to functional variation in fitness-related traits. In other cases, functional experiments on molecular function at certain loci may prompt population genetic tests to investigate whether the detected functional variation is of adaptive significance (Storz & Wheat, 2010). Here we present results from a study that followed the latter approach. Microsatellite loci that showed significant associations between allele lengths and gene expression levels in common sunflower (*Helianthus annuus* L.) were investigated to determine whether their population genetic structures were consistent with these loci playing a role in observed patterns of local adaption across a well-defined cline.

Previously, with an RNAseq approach, we identified 479 transcribed microsatellites in common sunflower where allele length significantly correlated with gene expression levels in a stepwise manner consistent with the predictions of the tuning knob model (hereafter referred to as eSTRs or significant expression short tandem repeats) (Ranathunge *et al*., 2020) (Figure 1a). These eSTRs are located in potential candidate genes for local adaptation, some of which have been linked to flowering time, disease resistance, and drought tolerance, among others. In the current study we test whether this functional variation detected in connection with the microsatellites is of adaptive significance with the use of polymorphism-based population genetic tests (Figure 1b).

**Figure 1.**
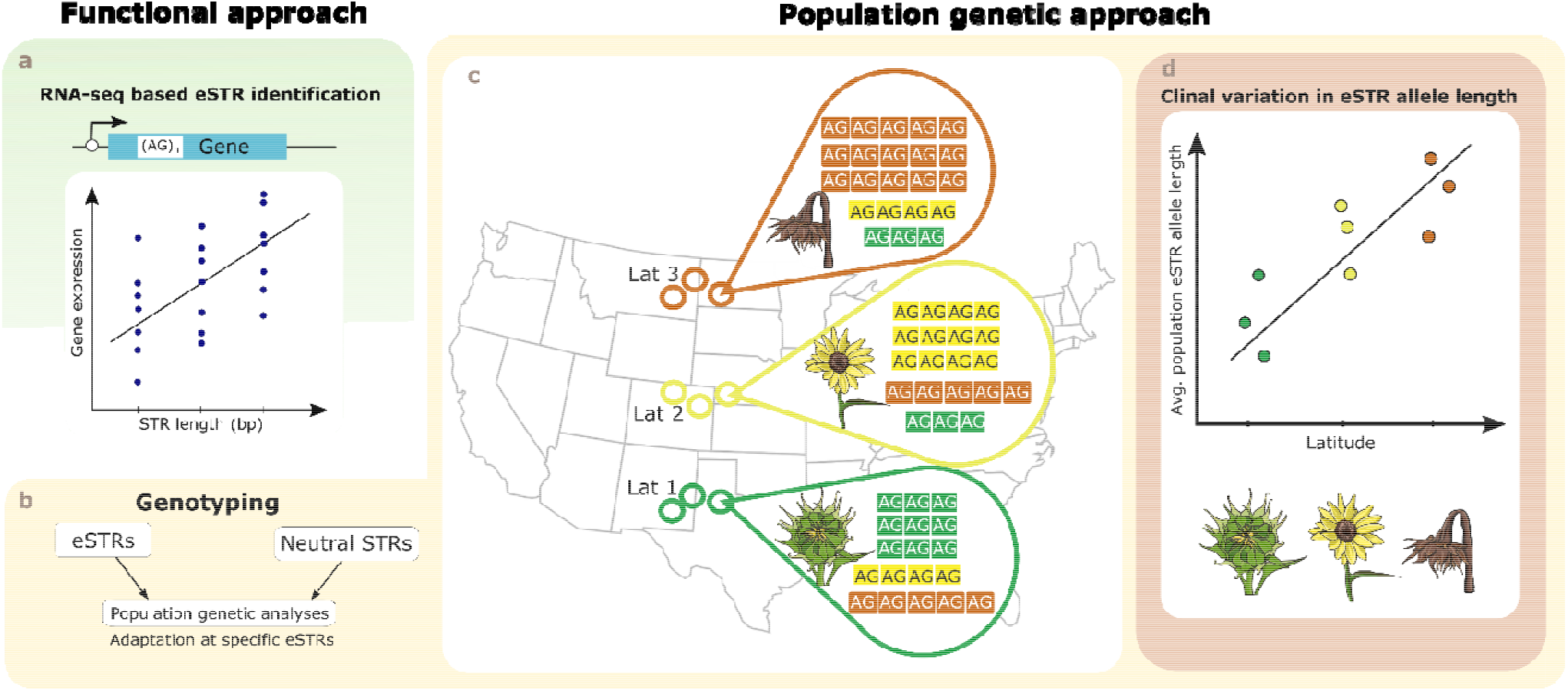
Integrating functional and population genetic approaches to infer adaptation at STRs. **a. Schematic diagram of eSTR identification using RNA-seq**. Previously, we identified 479 eSTRs where allele length significantly correlated with gene expression in sunflower using an RNA-seq approach (Ranathunge et al., 2020) **b. Population genetic analysis**. We genotyped 13 eSTRs previously identified with RNA-seq along with 19 neutral STRs and conducted population genetic analyses to test whether eSTR allele length is under selection. **c. Schematic diagram of a hypothetical model showing eSTR allele length variation in sunflower populations across a latitudinal cline**. Alle frequencies at a specific eSTR is shown here to be significantly different among sunflower populations across a well-defined cline where populations show continuous variation in flowering time. **d. Clinal variation in average population eSTR allele length**. Consistent with the predictions of the tuning knob model eSTRs show clinal patterns of variation in allele length with shorter or longer alleles being favored in climatic extremes.

As previously described, natural populations of common sunflower exhibit adaptations to a diverse range of habitats, and latitude is known to correlate with heritable variation in several traits such as flowering time (Blackman *et al*., 2011), seed oil content (Linder, 2000), and plant height (McAssey *et al*., 2016). Variation in transcribed microsatellite lengths and variation in gene expression levels of these loci has also previously been observed (Pramod et al., 2011, Pramod et al., 2012). In this study, we examined sunflower populations across a broad latitudinal transect running from Saskatchewan to Oklahoma, beyond the focal range from Kansas to Oklahoma considered in the previous RNAseq experiment that identified the eSTRs (Ranathunge et al, 2020). The inclusion of populations beyond the focal range also allowed us to test the hypothesis that shorter or longer allele lengths at eSTRs may be favored in more extreme environments (Figure 1c, d).

Here we assessed the strength of selection acting on a subset of eSTRs (13) compared to that acting on presumably neutral anonymous microsatellites (19). We compared the two sets of microsatellites on genetic diversity and allele frequency-based estimates under the hypothesis that eSTRs would show significantly higher or lower values for those estimates compared to neutral microsatellites which would indicate stronger selective pressures acting on the eSTRs. We conducted population structure analyses with the two different sets of microsatellites, and further examined the loci for selective sweeps with the test statistics *lnRV* and *lnRH* to provide a robust assessment of selection on eSTRs. Here we present an in-depth analysis of selective pressures on microsatellites that have shown evidence of functionality and demonstrate that gene expression variance influenced by transcribed microsatellites may contribute to the heritable variation underlying local adaptation.

## METHODS

### Plant material

For this study we selected natural populations of common sunflower from across the same latitudinal cline where others have demonstrated continuous variation in certain traits such as flowering time (Blackman *et al*., 2011, Ranathunge *et al*., 2018), seed oil content (Linder 2000), seed mass (Ranathunge *et al*., 2018), and plant height (McAssey *et al*., 2016, Ranathunge *et al*., 2018). Seeds from 17 natural populations of common sunflower transecting the latitudinal cline from Saskatchewan to Oklahoma were collected and grown in either laboratories or greenhouses (Figure 2; Supplementary Table S1). DNA was isolated from leaf tissue with Maxwell 16 tissue DNA purification kits (Promega, WI, USA), Qiagen DNeasy Plant Mini Kits (Valencia, CA), or the CTAB protocol (Murray & Thompson, 1980).

**Figure 2.**
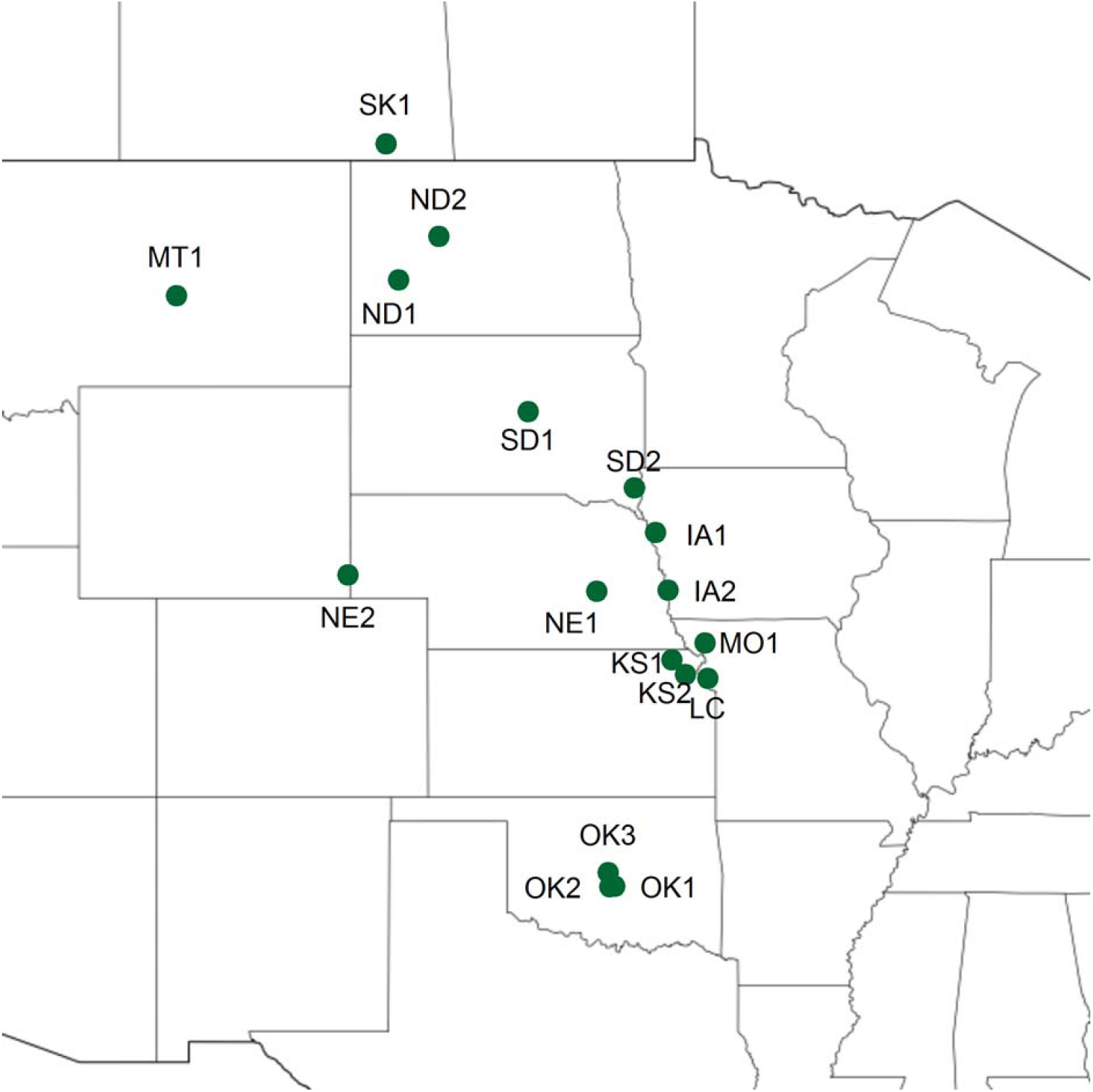
Geographical locations of the natural populations of common sunflower (*Helianthus annuus* L.) used in this study.

### Transcribed and anonymous microsatellite loci

A total of 13 eSTRs out of 479 identified by Ranathunge *et al*. (2020) were selected for this study (Table 1) to assess the strength of selection acting on functional transcribed microsatellites relative to that on anonymous microsatellites that are assumed to evolve neutrally. Microsatellite motif information and the putative functions of the 13 eSTR-containing genes identified by a BLASTX search against the *Helianthus annuus* protein sequence database (Ranathunge *et al*., 2020) are given in Table 1. The 19 anonymous microsatellites used in this study, and their PCR primers, were previously described by Tang *et al*., (2002) and Yu *et al*., (2002).

**Table 1.** Microsatellite repeat motif, location of the microsatellite within the gene and the putative functions of the eSTR-containing genes used in the study.

| eSTR-containing gene | Putative function | Microsatellite motif |
| --- | --- | --- |
| comp47993 | Dual specificity protein phosphatase DSP8 | TTCAA |
| comp43727 | Pentatricopeptide repeat (PPR) superfamily protein | TTG |
| comp25013 | ATP-dependent Clp protease proteolytic subunit 5, chloroplastic-like | GACGGT |
| comp47327 | Katanin p60 ATPase-containing subunit A1-like | A |
| comp47056 | Probable ADP-ribosylation factor GTPase-activating protein AGD14 isoform X1 | ATCA |
| comp36587 | Uncharacterized protein | GGTGAT |
| comp50288 | Receptor protein kinase-like protein ZAR1 | CA |
| comp41936 | ATP synthase delta chain, chloroplastic-like | ACCGCC |
| comp45709 | Cytochrome P450 86A22-like | GTGTTT |
| comp50462 | Probable pre-mRNA-splicing factor ATP-dependent RNA helicase DEAH5 | CGATCC |
| comp26672 | Protein CHUP1, chloroplastic-like | CCTTCT |
| comp45165 | Probable zinc metalloprotease EGY1, chloroplastic | ATA |
| comp25591 | Putative LSM domain-containing protein | AC |

### PCR and genotyping

Primers for the 13 eSTRs were designed for this study with Primer 3 development software (Untergasser *et al*., 2012) (Supplementary Table S2). Forward primers were either labeled with 6FAM, NED, PET, or VIC fluorescent dyes (Schuelke, 2000) or a M13 extension (CACGACGTTGTAAAACGAC) was added (Supplementary Table S2). PCR was performed in 10 µL volumes with ∼10 ng of DNA, 2 mM MgCl_2_, 30 mM tricine (pH 8.4) –KOH, 50mM KCl, 100 µM of each dNTP, 200 – 300 nM of reverse primer, 150 – 300 nM of forward primer, 200 nM of M13 forward primer, and 0.4 U of Taq DNA polymerase. We used touchdown – PCR protocols (Don *et al*., 1991) developed to reduce nonspecific primer binding and fragment amplification. The touchdown – PCR profiles included an initial denaturation step at 94 ^0^C for 5 min followed by 10 touchdown cycles at 94 ^0^C for 30 s, appropriate annealing temperature (touchdown annealing temperature = annealing temperature of the primer + 10 ^0^C) for 30 s, and another 30 s at 72 ^0^C. The annealing temperature decreases by 1 ^0^C in each successive touchdown cycle. The thermal cycle profiles for the remaining 25 cycles included 30 s at 94 ^0^C, 45 s at the appropriate annealing temperature for each primer (Supplementary Table S2), 30 s at 72 ^0^C, and a final elongation step of 7 min at 72 ^0^C. Fragment analyses were performed on ABI 3730 capillary sequencers (Applied Biosystems) at the Arizona State University DNA laboratory. LIZ 500 (GeneScan – 500 LIZ Size Standard – Applied Biosystems) was used as the size standard. Genotypes were visually scored using Peak Scanner version 1.0 and GeneMarker V2.6.7 (SoftGenetics, USA) (Supplementary Tables S3, S4).

### Genetic diversity and population structure

Genetic diversity at the 13 eSTR and 19 anonymous microsatellite loci was measured by estimating observed (H_O_) and expected heterozygosity (H_E_) values according to Nei (1978) with Arlequin v. 3.5.1.3 (Excoffier & Lischer, 2010). Pairwise F_ST_ values (Weir & Cockerham, 1984) were estimated for each microsatellite locus with Arlequin 3.5.1.3 (Excoffier & Lischer, 2010). The number of alleles per locus (A) and mean number of alleles per locus or allelic diversity (AD) for each eSTR and anonymous microsatellite locus were measured with GenAlEx v. 6.503 (Peakall & Smouse, 2012). We conducted Wilcoxon rank sum tests to compare H_E_, F_ST_, A and AD estimates between the two types of microsatellites. Mantel tests were used to assess whether correlation between geographic and genetic distance matrices at the two types of loci was significant (Mantel, 1967). Genetic distance matrices were computed using Slatkin’s pairwise F_ST_ estimates (Slatkin, 1987) obtained from Arlequin (Excoffier & Lischer, 2010). Mantel tests for the two types of loci were performed between pairwise linearized genetic distance and log transformed geographic distance matrices with the program IBD (Isolation by Distance) (Bohonak, 2002).

To test the hypothesis that the number of distinct populations identified within the data set by the two types of loci could be different due to differences in the evolutionary processes acting on them, we conducted a Bayesian-based individual assessment test with the program STRUCTURE v. 2.3.4 (Pritchard *et al*., 2000) for eSTRs and anonymous microsatellites separately. The program attempts to assign individuals to K number of distinct populations based on the posterior probability that a set of genotypes were sampled from a model allelic distribution. The model used in the program simulates an admixture of individual ancestry with correlated allele frequencies. No prior information on populations was included in the model. We set the length of the burn-in to 200,000, and a total of 800,000 MCMC (Markov Chain Monte Carlo) iterations were calculated. A total number of 20 iterations were conducted for each value of K (from 2 to 20). Based on the posterior probability values calculated for each value of K, we estimated the most likely number of clusters that best explains the partitioning of genetic variation at the two types of microsatellites with the “ Evanno” method (Evanno *et al*., 2005) implemented by the program Structure Harvester (Earl & vonHoldt, 2012).

### Loci under selection

Tests for evidence of selection were conducted on the two types of microsatellites with *lnRV* and *lnRH* test statistics. *lnRV* is estimated based on the variance in number of repeats (Harr *et al*., 2002; Schlötterer, 2002), and *lnRH* based on the expected heterozygosity (Schlötterer & Dieringer, 2005). *lnRV* and *lnRH* are two powerful test statistics used to identify loci that have undergone strong selection. Variance in the number of repeats (V) or heterozygosity (H) at a microsatellite locus in a given population can be considered a function of effective population size (N_e_) and microsatellite mutation rate (μ) and estimating ratios of variances in repeat numbers and heterozygosity removes locus-specific effects which makes *lnRV* and *lnRH* measures of variation in N_e_. The test statistics, *lnRV* and *lnRH* can be calculated for population pairs as follows,

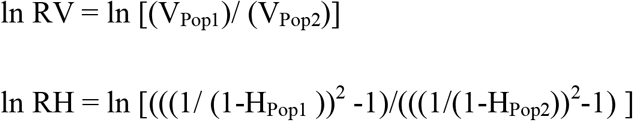

Under neutrality, values for both statistics are considered to follow a normal distribution pattern with values falling outside of −1.96 to 1.96 interval corresponding to outliers caused by selection (p < 0.05). When both statistics are used together, the number of false positives (Type I error) is reduced by a factor of three (Schlötterer & Dieringer, 2005).

To calculate *lnRV* and *lnRH* statistics, we first estimated variance in the number of repeats (V) and expected heterozygosity (H) for each microsatellite locus in each population with Microsatellite Analyzer (Dieringer & Schlötterer, 2003). Pairwise estimates for *lnRV* and *lnRH* were calculated for each population pair (136 comparisons in total among the 17 populations) for each microsatellite locus. If a population was monomorphic for a certain locus, we added a single allele that differed in one repeat unit to avoid heterozygosity being calculated as zero, an approach that has previously been successfully used (Schlötterer & Dieringer, 2005; Vasemägi *et al*., 2005; Coscia *et al*., 2012). The calculated *lnRV* and *lnRH* values were standardized by the mean and standard deviation according to Kauer *et al*. (2003). We conducted Chi-squared analyses to test whether higher numbers of outliers for *lnRV* and *lnRH* identified at eSTRs compared to anonymous microsatellite loci were statistically significant, a finding that would be consistent with selection acting on locally adaptive alleles at the eSTRs.

### Latitudinal variation in microsatellite allele length

The eSTRs used in this study have shown significant allele length effects on gene expression in common sunflower across populations from a narrow latitudinal range between Kansas (KS) and Oklahoma (OK) (Ranathunge *et al*., 2020). Given that microsatellite allele length has been linked to gene expression across a narrow latitudinal range, we hypothesized that this variation may be functional and locally adaptive. Hence, we predicted that a significant portion of the variation in allele length at the eSTRs could be explained in relation to environmental conditions associated with latitudes. We further predicted that longer or shorter microsatellite alleles would be favored in populations located in more extreme latitudes beyond the focal populations sampled in our previous study (Ranathunge *et al*., 2020). This hypothesis is consistent with the predictions of the tuning knob model which suggests that variation in microsatellite allele length can have stepwise effects on phenotypes (Kashi *et al*., 1997; King *et al*., 1997; Trifonov, 2004). To test whether populations from more extreme latitudes harbor longer or shorter microsatellite alleles relative to the focal populations from Kansas and Oklahoma, we conducted linear regression analyses for both eSTRs and anonymous microsatellites with population mean microsatellite allele length at a locus as the response variable and the latitudinal location of the populations as the continuous predictor variable. The proportion of allele length variation explained by latitude was assessed by means of ANOVA using R statistical software (R Core Team, 2017).

## RESULTS

### Comparison of genetic diversity measures in eSTR and anonymous microsatellite loci

The 13 eSTRs used in this study consisted of one mononucleotide, two dinucleotides, two trinucleotides, one tetranucleotide, one pentanucleotide, and six hexanucleotides repeats (Table 1). The putative functions associated with the 13 eSTR-containing genes indicated their potential role in local adaptation of *H. annuus* (Table 1). To assess the selective pressures acting on the eSTRs, we compared genetic diversity estimates obtained for the 13 eSTRs with that observed at the 19 anonymous microsatellite loci used in this study.

Observed heterozygosity (H_O_) at the eSTRs ranged from 0.308 to 0.89. H_O_ at the anonymous microsatellites ranged between 0.223 and 0.715 (Table 2). Expected heterozygosity (H_E_) ranged from 0.26 to 0.794 across the 13 eSTRs and from 0.689 to 0.87 across the 19 anonymous microsatellite loci (Table 2). The mean H_E_ estimates were 0.672 and 0.781 at the eSTRs and the anonymous microsatellites, respectively. Loci comp36587 and comp47327 showed the lowest and the highest H_E_ among the eSTRs, respectively (Table 2). H_E_ values at the eSTRs were significantly lower compared to that at anonymous microsatellite loci (Wilcoxon rank sum test, p = 0.002241) (Figure 3a). Allelic richness (A) and allelic diversity (mean number of alleles) (AD) at eSTRs and anonymous microsatellites also differed significantly between the two types of microsatellites. Allelic richness (A) ranged between four and 16 for eSTRs and between eight and 25 for the anonymous microsatellite loci (Table 2). eSTRs, comp36587 and comp45165 showed the lowest and the highest A (Table 2), respectively. The mean allelic richness across eSTRs and anonymous microsatellites were 9.923 and 17.158, respectively. The estimates for A were significantly lower at the eSTRs compared to that at the anonymous microsatellites (Wilcoxon rank sum test, p = 0.001392) (Figure 3d). A similar trend was observed in estimates for AD between the two types of loci. The estimates for AD ranged from 2.647 to 9.529 in the eSTRs and from 5.294 to 13.588 in the anonymous microsatellites (Table 2). The lowest AD value among the eSTRs was observed in comp36587 and the highest in comp50288 (Table 2). The mean AD across eSTRs and anonymous microsatellites were 6.443 and 9.607, respectively. AD was significantly low in the eSTRs compared to anonymous microsatellites (Wilcoxon rank sum test, p = 0.002594) (Figure 3c). Estimates for F_ST_ ranged from 0.048 to 0.108 in the eSTRs and from 0.043 to 0.112 in the anonymous microsatellites. (Table 2). Locus comp50288 showed the highest estimate for F_ST_ among the eSTRs and the lowest estimate was observed at comp25013 (Table 2). The mean F_ST_ at the eSTRs was 0.071 and 0.072 at the anonymous microsatellite loci. The estimates for F_ST_ were not significantly different between the two types of microsatellites (Wilcoxon rank sum test, p = 0.8501) (Figure 3b).

**Table 2.** Allelic diversity (AD), allelic richness (A), allele size range, observed heterozygosity (H_O_), expected heterozygosity (H_E_) and pairwise F_ST_ estimates at transcribed (eSTRs) and anonymous microsatellite loci.

| Locus | Allele size range (bp) | AD | A | $H_O$ | $H_E$ | $F_{ST}$ |
| --- | --- | --- | --- | --- | --- | --- |
| comp47993 | 221 – 226 | 4.118 | 6 | 0.668 | 0.619 | 0.049 |
| comp43727 | 191 – 221 | 7.765 | 11 | 0.732 | 0.661 | 0.096 |
| comp25013 | 184 – 208 | 4.471 | 5 | 0.692 | 0.635 | 0.048 |
| comp47327 | 135 – 167 | 8.529 | 12 | 0.852 | 0.794 | 0.051 |
| comp47056 | 215 – 253 | 7.471 | 10 | 0.859 | 0.747 | 0.090 |
| comp36587 | 227 – 239 | 2.647 | 4 | 0.308 | 0.260 | 0.069 |
| comp50288 | 238 – 266 | 9.529 | 15 | 0.890 | 0.724 | 0.108 |
| comp41936 | 294 – 324 | 3.941 | 6 | 0.664 | 0.603 | 0.074 |
| comp45709 | 320 – 350 | 4.588 | 6 | 0.738 | 0.686 | 0.054 |
| comp50462 | 188 – 218 | 7.000 | 11 | 0.829 | 0.760 | 0.061 |
| comp26672 | 241 – 271 | 6.176 | 11 | 0.824 | 0.754 | 0.056 |
| comp45165 | 184 – 232 | 8.353 | 16 | 0.878 | 0.763 | 0.084 |
| comp25591 | 162 – 206 | 9.176 | 16 | 0.845 | 0.726 | 0.089 |
| ORS878 | 207 – 247 | 10.647 | 21 | 0.378 | 0.800 | 0.079 |
| ORS483 | 272 – 294 | 7.706 | 12 | 0.592 | 0.734 | 0.109 |
| ORS420 | 144 – 164 | 6.706 | 11 | 0.669 | 0.753 | 0.043 |
| ORS1161 | 235 – 287 | 13.588 | 25 | 0.514 | 0.870 | 0.047 |
| ORS456 | 331 – 345 | 5.294 | 8 | 0.492 | 0.689 | 0.066 |
| ORS299 | 272 – 288 | 6.529 | 9 | 0.406 | 0.700 | 0.062 |
| ORS1067 | 190 – 226 | 12 | 19 | 0.645 | 0.848 | 0.064 |
| ORS1088 | 205 – 306 | 12.941 | 25 | 0.454 | 0.862 | 0.050 |
| ORS1108 | 173 – 209 | 10.294 | 16 | 0.565 | 0.787 | 0.086 |
| ORS1144 | 137 – 169 | 9.471 | 17 | 0.475 | 0.784 | 0.091 |
| ORS1193 | 306 – 332 | 6.882 | 10 | 0.349 | 0.736 | 0.112 |
| ORS380 | 400 – 456 | 11.941 | 22 | 0.587 | 0.846 | 0.058 |
| ORS437 | 343 – 383 | 8.765 | 16 | 0.366 | 0.702 | 0.075 |
| ORS608 | 271 – 329 | 8.471 | 23 | 0.223 | 0.76943 | 0.07026 |
| ORS613 | 225 – 259 | 11.353 | 18 | 0.524 | 0.83484 | 0.0798 |
| ORS691 | 453 – 499 | 12.412 | 22 | 0.414 | 0.83552 | 0.07357 |
| ORS815 | 189 – 243 | 10.529 | 18 | 0.715 | 0.7737 | 0.06973 |
| ORS853 | 185 – 247 | 10.235 | 23 | 0.443 | 0.80938 | 0.07117 |
| ORS857 | 220 – 242 | 6.765 | 11 | 0.438 | 0.71011 | 0.06589 |

**Figure 3.**
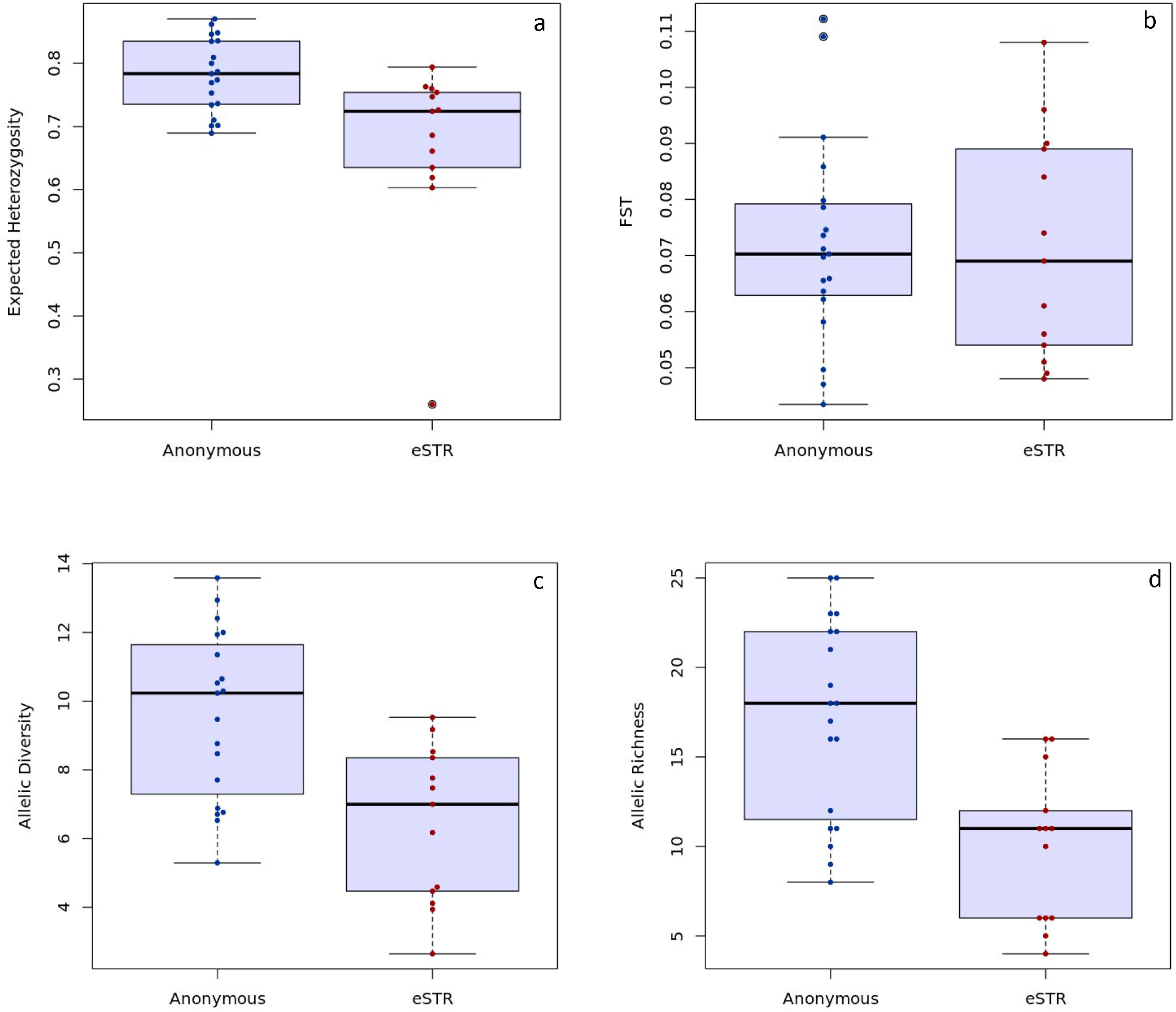
Expected Heterozygosity (H_E_) (a), population genetic differentiation (b), allelic diversity (c), and allelic richness (d) estimates across 17 natural populations of common sunflower (*Helianthus annuus* L.) at 19 anonymous and 13 transcribed microsatellite loci (eSTRs) used in the study.

Isolation by distance analyses based on Mantel tests showed significant positive correlation between genetic and log transformed geographic distance matrices for both eSTRs (R = 0.3824, p < 0.001), and anonymous microsatellites (R = 0.3770, p < 0.001).

### Population structure

STRUCTURE analyses conducted separately for the two types of microsatellites indicated distinct patterns of individual assignment to clusters. The DeltaK method (Evanno *et al*., 2005) implemented with the Structure Harvester program (Earl & von Holdt, 2012) determined the most likely number of clusters that explained the genetic variation at the two types of microsatellites. The most likely number of clusters supported when using anonymous microsatellites was K =2. (Supplementary Figure S1a; Supplementary Figure S2a). At the eSTRs, the most likely number of clusters that best explained the partitioning of genetic variation across the 672 individuals was K = 5 (Supplementary Figure S1b;; Supplementary Figure S2b).

This finding is consistent with elevated levels of linkage disequilibrium among eSTRs relative to that observed among anonymous microsatellites. At the anonymous microsatellites, the clustering pattern suggested a north-south division in the populations with the transition zone close to populations from Missouri and Kansas (Supplementary Figure S1a). At the eSTRs, populations ND1 and MT1 formed a distinct cluster from the remaining 15 populations, and so did population NE2 (Supplementary Figure S1b). The north-south division observed when the structure analysis was conducted with anonymous microsatellites was also detected with eSTRs, although not prominently so.

### Loci under selection

Significant *lnRV* values based on the variance in the number of repeats were detected in all 13 eSTRs and 18 of the19 anonymous microsatellites (p < 0.05) (Supplementary Table S5). A total of 141 *lnRV* outliers were detected across the eSTRs and 167 across the anonymous microsatellites. The number of *lnRV* outliers per locus ranged from two to 16 for eSTRs and from five to 14 for anonymous microsatellites based on all possible population pair comparisons (136) for each microsatellite locus (Supplementary Table S5). Collectively, there was no significant difference in the number of *lnRV* outliers produced by the eSTRs compared to that produced by the anonymous microsatellites (Chi-squared test, p = 0.06425). The total number of *lnRH* outliers identified across the 13 eSTRs and the 19 anonymous microsatellites was 123 and 138, respectively. The number of significant *lnRH* values per locus ranged from four to 15 for eSTRs and from five to 14 for anonymous microsatellite loci (Supplementary Table S5). The number of *lnRH* outliers detected in eSTRs was significantly higher than that observed across anonymous microsatellites (Chi-squared test, p = 0.03229). As suggested by Schlötterer & Dieringer (2005), we regressed *lnRV* against *lnRH* for each population comparison-a conservative approach that should reduce the number of false positives by a factor of three. The number of population comparisons with significant values for both *lnRV* and *lnRH* ranged from seven to 11 across 11 of the 13 eSTRs and one to five across 15 of the 19 anonymous microsatellites (Figure 4; Supplementary Table S5). Collectively, eSTRs produced significantly more outliers (58) with significant values for both *lnRV* and *lnRH* than did anonymous microsatellites where only 32 such outliers were identified (Chi-squared test, p =0.000005601) (Figure 4; Supplementary Table S5.

**Figure 4.**
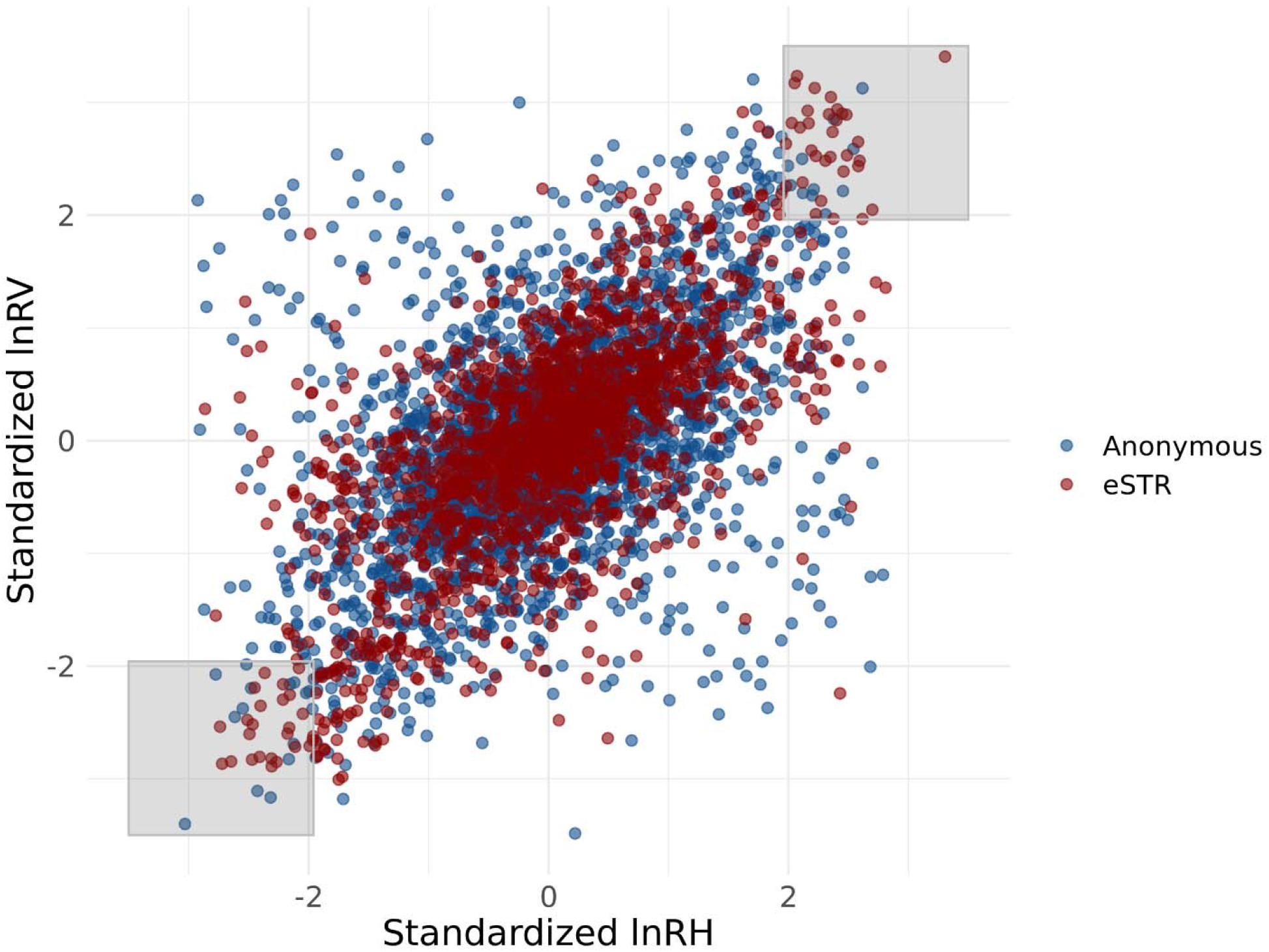
The relationship between standardized *lnRV* and standardized *lnRH*. Data points represent *lnRV* and *lnRH* values calculated for each sunflower (*Helianthus annuus* L.) population pair at each microsatellite locus. The gray boxes indicate the regions where data points are outliers for both *lnRV* and *lnRH* (p < 0.05).

### Microsatellite allele length variation across latitudes

Based on the patterns of correlation observed between eSTR allele length and gene expression in sunflower populations across a narrow latitudinal range (Ranathunge *et al*., 2020), we predicted clinal variation in eSTR allele lengths across a broader latitudinal range. Such patterns in eSTR allele length coupled with the results from previous functional analyses (Ranathunge et al., 2020), would provide further evidence in support of a role for transcribed microsatellites in facilitating local adaptation. To test whether sunflower populations from more extreme latitudes harbor shorter or longer eSTR allele lengths compared to the focal populations from Kansas and Oklahoma, we calculated population mean allele length for each eSTR locus and anonymous microsatellite locus and conducted linear regression analyses between population mean allele length and latitude. The proportion of allele length variation explained by latitude varied from 0.15% to 50.41% across the eSTRs. Six of the 13 eSTRs showed a negative linear relationship between mean population allele length and latitude (Supplementary Table S6) which shows that shorter allele lengths are more common at higher latitudes at these six eSTRs. The opposite trend (positive linear relationship) was observed at the remaining seven eSTRs with northern populations harboring longer allele lengths compared to the south. For four of the 13 eSTRs the correlation between mean population allele length and latitude was statistically significant. comp41936 (p < 0.05) and comp50288 (p < 0.01) showed a significant positive linear relationship between eSTR mean population allele length and latitude (Figure. 5a, b) while comp25591 (p < 0.05) and comp47993 (p < 0.01) showed significant negative linear relationship (Supplementary Table S6) (Figure 5b, d). In comparison, two of the 19 anonymous microsatellite loci, ORS 815 and ORS857, showed statistically significant correlation between mean population allele length and latitude. ORS857 showed a significant negative linear relationship (p < 0.05) while ORS815 showed a significant positive linear relationship (p < 0.01) between population mean microsatellite allele length and latitude (Figure 5e, f). The proportion of allele length variation explained by latitude varied from 0.02% to 50.4% across the 19 anonymous microsatellites.

**Figure 5.**
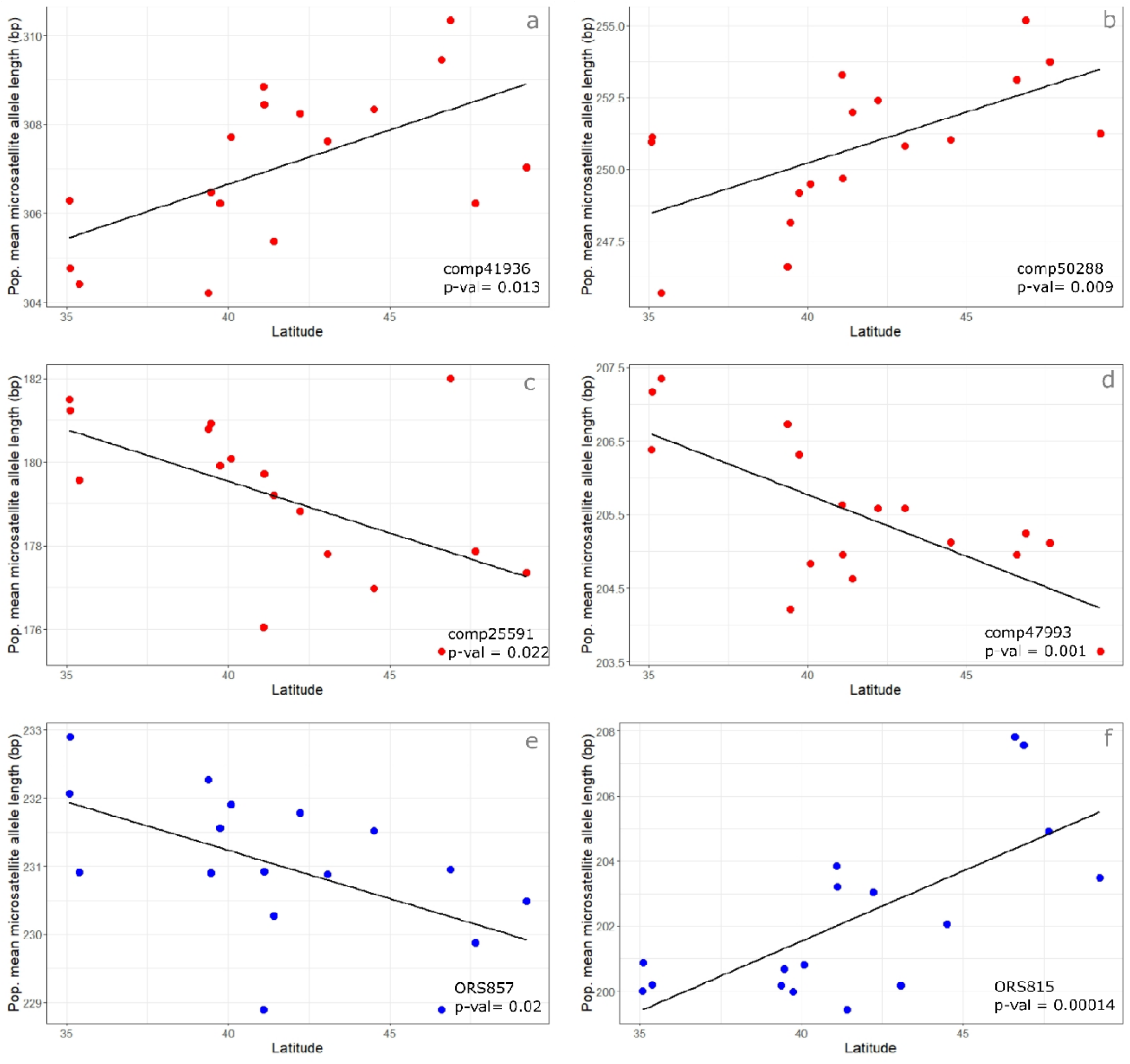
Population mean microsatellite allele length plotted as a function of latitude for eSTR loci (a) comp41936, (b) comp50288, (c) comp25591 (d) comp47993, and anonymous loci, (e) ORS857 and (f) ORS815 across 17 common sunflower (*Helianthus annuus* L.) populations.

## DISCUSSION

Although long considered neutrally evolving regions in the genome, research now implies that microsatellites are capable of functioning as advantageous mutators (Kashi *et al*., 1997; King *et al*., 1997; Trifonov, 2004). With an RNA-Seq approach, we previously identified transcribed microsatellites that influence gene expression in a stepwise manner (eSTRs) in natural populations of common sunflower across a narrow latitudinal range between Kansas and Oklahoma (Ranathunge *et al*., 2020). Designating these eSTRs as candidates for rapid adaptive evolution based on their measurable effects on functional variation alone is insufficient. Here we tested whether these eSTRs are under selection, and whether these eSTRs show clinal patterns in genetic variation consistent with the predictions of the tuning-knob model which could provide further support to their role as advantageous mutators.

Collectively, genetic diversity estimates obtained from the eSTRs and anonymous microsatellites are consistent with purifying selection having a greater influence on eSTRs compared to that acting on anonymous microsatellites. Estimates for expected heterozygosity (H_E_), allelic richness (A), and allelic diversity (AD) were significantly low at the 13 eSTRs in comparison to that at anonymous microsatellites (Figure. 3a, c, d; Table 2). Similar trends have been reported in several studies that have compared microsatellites located within transcribed regions (EST-SSRs) to anonymous microsatellites (genomic SSRs) suggesting that EST-SSRs are generally less polymorphic than genomic microsatellites (Cho *et al*., 2000; Leigh *et al*., 2003; Eujayl *et al*., 2004; Varshney *et al*., 2005; Yatabe *et al*., 2007). Low levels of polymorphism detected at transcribed microsatellites have been typically attributed to functional and evolutionary constraints that limit expansion of specific repeats within structurally important regions (Dokholyan *et al*., 1997; Metzgar *et al*., 2000). However, other factors likely contribute to lower relative diversity at eSTRs and cannot on their own be viewed as strong evidence for purifying selection in this case. For one, genetic variation typically observed at genic microsatellites may not necessarily indicate selection acting on the microsatellite itself, rather it may reflect linkage disequilibrium between the microsatellite and a region, or regions, under selection. Given that all of the eSTRs identified here are located within genes heightens this likelihood. Lower diversity at the eSTRs may also reflect differences in mutation rates. The 13 eSTRs have a greater average motif length (Table 1) than do the anonymous microsatellites examined in this study and STR mutation rates are negatively correlated with the length of these repeat units (Kelkar *et al*., 2009).

Theoretically, the proportion of allele frequency variation attributed to population differences is determined by selection (Lewontin & Krakauer, 1973). This proportion of genetic variation identified as F_ST_ (Wright, 1951) is considered to vary significantly from neutrally evolving loci due to selection. In this study, we did not detect any significant difference in mean F_ST_ values at eSTRs compared to anonymous microsatellites which suggests that in general, eSTRs are not under stronger selective pressures compared to anonymous microsatellites. However, there is greater relative variance in F_ST_ at the eSTRs compared to anonymous microsatellites (Figure. 3b) which is consistent with the hypothesis that different patterns of selection may act on different eSTR loci (Table 2).

Inferences based on F_ST_ alone can be error prone where microsatellites are considered due to high mutation rates at microsatellites (Hedrick, 1999; Balloux et al., 2000; Putman & Carbone, 2014), differences in mutation rates among loci (Storz *et al*., 2004), or locus-specific phenomena linked to selection (Schlötterer & Dieringer, 2005). To provide support for patterns of selection detected via F_ST_ based tests, we used a second, more robust approach with *lnRV* and *lnRH* test statistics that accounts for some of these problematic features of microsatellites such as locus specific mutation rates, for identifying loci under selection. The two statistics *lnRV* and *lnRH* identify loci under selection based on variability in the number of repeats and heterozygosity, respectively (Schlötterer, 2002; Schlötterer & Dieringer, 2005). Our results from the *lnRV* and *lnRH* based analyses revealed that the number of outliers for *lnRH* at eSTRs significantly outnumbered those at anonymous microsatellites (Figure 4; Supplementary Table S5). Furthermore, outliers for both *lnRV* and *lnRH* combined were also significantly higher in the eSTRs compared to anonymous microsatellites. Combined with the greater variance in F_ST_ observed at the eSTRs, these outliers for *lnRV* and *lnRH* suggest that different eSTRs maybe under different rates of selection compared to anonymous microsatellites. Further, our results from the *lnRV* and *lnRH* analysis suggest that eSTRs are likely to have undergone recent local selective sweeps whereby different allele lengths may be favored in different populations. We detected fixed alleles at three eSTRs (comp50288, comp36587, and comp25591) across two populations (ND1, NE2) that suggest a putative sweep at the population level. At comp 36587, both ND1 and NE2 populations were monomorphic for allele size 233 bp. Loci comp50288 and comp 25591 each had a fixed allele in populations NE2 (252 bp) and ND1 (182 bp), respectively. Similar selective sweeps have been previously reported for expressed microsatellites at inter and intra species levels in the genus *Helianthus* across North America (Gross *et al*., 2007; Kane & Rieseberg, 2007; Chapman *et al*., 2008; Kane *et al*., 2009). In all such microsatellite-based studies in sunflower that reported selective sweeps, authors have emphasized that the extreme *lnRV* and *lnRH* estimates do not indicate that the microsatellites themselves are under selection rather that they could be tightly linked to loci under selection. In the absence of functional experimental data to suggest otherwise, this rationale that microsatellites could be linked to adjacent candidate loci seems plausible. However, it is also interesting to note that linkage disequilibrium in wild sunflower genomes is considered to be extremely low which suggests that regions that have undergone selective sweeps are likely to be small (Liu & Burke, 2006; Kolkman *et al*., 2007).

Our population structure results are consistent with previous studies and suggest that the anonymous microsatellite loci used in this study are effective at detecting patterns of genetic differentiation previously inferred to reflect neutral rates of gene flow across this cline (Mandel *et al*., 2011; McAssey *et al*., 2016). Comparatively, partitioning of genetic variation at eSTRs was different from that at anonymous microsatellites, which may indicate distinct selective pressures acting on the eSTRs. Pramod *et al*., (2012) also reported different patterns of selection acting on transcribed microsatellites compared to anonymous microsatellites in common sunflower across populations from a narrower latitudinal range to that of the current study.

In line with the signatures of directional selection detected at the eSTRs, we predicted that eSTR mean microsatellite allele lengths could be tightly linked to the location of the population. This prediction follows directly from the tuning-knob hypothesis and assumes that clinal patterns of variation in several heritable traits in sunflower populations across this well-defined cline could be linked to variation in eSTR length We detected statistically significant relationships between population mean allele length and latitude at four of the 13 eSTRs (Figure 5; Supplementary Table S6). comp50288 and comp41936 showed strong positive correlation while comp49773 and comp25591 showed a negative correlation between population mean allele length and latitude. In comparison, two of the 19 anonymous microsatellites showed significant correlation between latitude and allele length (Figure 5). Collectively, our results show that it is more likely that shorter or longer alleles at the eSTRs may be favored in extreme latitudes. The strong correlation between eSTR allele length and latitude observed in this study (Figure 5) coupled with the results from the previous RNA-seq based functional experiments (Ranathunge et al. 2020) strengthens the argument that eSTRs themselves could be under selection. Similar patterns of microsatellite allele length variation across latitudes have been detected in previous studies with particular focus on genes related to circadian rhythm (Johnsen *et al*., 2007; Michael *et al*., 2007; Lemay & Russello, 2014). Note that many adaptive gradients likely correlate poorly with latitude. Hence, tuning knobs at those loci would likely go undetected using this approach.

In this study we assessed the strength of selection acting on microsatellites that have shown evidence for functionality in sunflower. Collectively, the results from the genetic diversity estimates, *lnRV* and *lnRH* based outlier tests, and latitudinal variation in mean microsatellite allele length suggest that eSTRs could be under directional selection. Our study provides compelling evidence to support the adaptive potential of transcribed microsatellites that contribute to variation in gene expression in common sunflower and demonstrate that natural populations may be able to generate heritable genetic variation through mutation at rates far greater than previously assumed.

## SUPPORTING INFORMATION

Figure S1. Delta K values calculated according to Evanno’s method showing the number of clusters best supported at (a) anonymous and (b) transcribed microsatellites (eSTRs). The number of clusters best supported at anonymous and transcribed microsatellite loci (eSTRs) was K = 2 and K = 5, respectively.

Figure S2. Population genetic structure of 17 natural populations of common sunflower (*Helianthus annuus* L.) at (a) anonymous and (b) transcribed microsatellite loci (eSTRs). The number of clusters (K) best supported at anonymous microsatellites (a) and eSTRs (b) was K = 2 and K = 5, respectively.

Supplementary Table S1. List of populations used in this study and their geographical locations

Supplementary Table S2. List of primers designed for the 13 eSTRs used in this study

Supplementary Table S3. Genotype data from the 19 anonymous microsatellite loci used in the study

Supplementary Table S4. Genotype data from the 13 transcribed microsatellite loci (eSTRs) used in the study

Supplementary Table S5. Standardized *lnRV* and *lnRH* estimates for eSTRs and anonymous microsatellites

Supplementary Table S6. Regression models estimating the effect of latitude on population mean eSTR allele length

